# A comparative metagenomic analysis of antibiotic resistance in the fecal microbiome of pigs raised with and without antibiotics

**DOI:** 10.1101/2025.01.20.633852

**Authors:** Stephanie Pillay, Thomas Abeel

## Abstract

**Purpose:** The use of antibiotics in swine production has raised concerns about its impact on animal health, microbial ecology, and the emergence of antibiotic resistance. The gut microbiome, a critical component of animal health, is susceptible to disruptions caused by antibiotic use. The fecal microbiome can provide a snapshot of changes in the gut microbiome and provide information on the spread of antibiotic resistance from pig feces to the environment.

**Methods:** We investigated the differences between the fecal microbiome of pigs raised with (AB+) and without (AB-) antibiotics to assess how antibiotic exposure influenced the microbial community, antibiotic resistance genes (ARGs), and mobile genetic elements (MGEs).

**Results:** Our findings revealed that the use of antibiotics combined with study-specific conditions alters the fecal microbiome. Antibiotic usage in pigs explains 20% of the variation in the diversity of bacterial species detected between AB+ and AB- pigs. Similarly, variations in ARGs were observed in the two groups. The RND efflux pump gene family was abundant in both groups. The OXA *β*-lactamase gene and associated plasmids were more prevalent in the AB- group. We also identified integrons linked to chloramphenicol resistance and transposon-associated integrative elements that can facilitate the spread of antibiotic resistance genes within and between bacterial species.

**Conclusion:** These findings highlight that antibiotics influence both the microbial community and the antibiotic resistance gene repertoire, regardless of study-specific or environmental conditions. This emphasises the potential for fecal matter to contribute to the spread of antibiotic resistance in the environment, underscoring the need for more research to fully understand these dynamics in agricultural settings.

## 1 Introduction

Antimicrobial resistance (AMR), resulting from excessive antibiotic use in humans, animals, and the environment, poses a significant global health risk. By 2030, antibiotic usage is projected to increase by 67%, with over half attributed to animals raised for food production [1, 2].

Antibiotics have been integral to the growth and development of food animals for more than half a century, particularly in swine production. Antibiotics are routinely used to treat, control and prevent diseases, and to increase productivity. Sub-therapeutic doses are used to control symptomatic infections between animals in close contact, to prevent disease at points of high risk, e.g. when the animal is stressed during extreme weather, post-vaccination or moving [3, 4]. This practice has turned the food animal industry into a hotspot for antibiotic resistant dissemination [2].

Among various food animals (cattle, poultry, and pigs), the pig industry is the most prolific user of antibiotics [5]. In pigs, antibiotics are given to whole groups by mixing antibiotics into feed or adding antibiotic powder or solution into drinking water [3]. Previous studies have shown that antibiotics, such as beta-lactams, tetracyclines, sulfonamides, lincosamides, macrolides, and quinolones, are typically used in swine production [6]. However, the choice of antibiotics is influenced by age-specific diseases, common pathogens, market availability, and cost [7]. Farm management practices also play a role in antibiotic use. Larger farms, due to their higher risk of pathogen transmission within herds, are more likely to use medicated feed compared to smaller farms. However, it has been proposed that while costly, good farm bio-security practices and vaccine administration reduce disease incidence and antibiotic use [8].

In recent years, the connection between antibiotic use in livestock and the emergence and spread of antibiotic resistance has been established. This has led to the European Union banning the “non-therapeutic” use of antimicrobials as growth promoters since 2006. Restrictions on the use of antibiotics followed in the United States since 2017 and China since 2020 [9]. Unfortunately, countries with large livestock production, such as China and Brazil, still remain the largest producer and consumer of antibiotics with 52% being used in animal production [7, 9]. The primary concern is that the antibiotics used in livestock sectors are also critically important for human health, including cephalosporins and fluoroquinolones. This can lead to bacteria becoming resistant to the last line of defence antibiotics and causing untreatable infections in humans and animals [10].

The gut microbiome of pigs plays a significant role in nutrition, metabolism and health. Alteration of the gut microbiome using probiotics or vaccines can prevent diseases leading to feeding efficiency, resistance to diarrhoea, meat quality, and immune function [11, 12]. Antibiotic usage influences the gut microbiome of pigs amongst other factors such as age, host genes, gender and the castration of male pigs. The gut microbiome harbours a diverse microbial community which supports the health of the animal, helping to exclude pathogens [9]. A highly diverse gut microbiome can become less diverse after antibiotic treatment causing lasting changes in the composition of the microbiome of pigs promoting antibiotic resistant bacteria and increasing the prevalence of antibiotic resistance genes (ARGs) [9]. Antibiotic resistant bacteria and ARGs can be excreted with fecal matter as a significant amount (30-80%) of antibiotics are excreted due to incomplete metabolism. Animal feces are commonly used as manure in agricultural practices in surrounding farms which can promote the spread of antibiotic resistance. Additionally, unconsumed feed, containing these antibiotics, can contaminate soil directly or indirectly affecting aquatic and farming systems through run-off [1].

Since it has been assumed that the fecal microbiome is a subset of the gut microbiome in pigs, it can be used to understand the effects of antibiotic usage on the gut microbiome, monitor the spread of antibiotic resistance to other environments and mitigate the risks by influencing farm policies [13]. We aim to use publicly available metagenomic data obtained from the fecal microbiome of pigs raised with and without antibiotics to identify and characterise the differences in the microbial community, ARGs and mobile genetic elements. Through this, we aim to establish the impact of antibiotics on the gut microbiome of pigs to broaden our understanding of antimicrobial resistance in food animals.

## Methods and Materials

### 1.1 Data Sources

103 publicly available metagenomic sequencing sets were obtained from studies that raised pigs with (AB+) and without antibiotics (AB-) (Table 1). Studies that had pigs raised without antibiotics and those that received antibiotics for illness only were categorised as “without antibiotics” (n = 56). Pigs raised with antibiotics were categorised as such (n = 47). Data was obtained from NCBI SRA (https://www.ncbi.nlm.nih.gov/sra) and the EBI MGNIFY database (https://www.ebi.ac.uk/metagenomics). A combination of keywords was used on NCBI SRA, these included “pigs”, “sus scrofra”, “feces”, “metagenomes” and “fecal”. On the EBI MGNIFY database, keywords included: biome = “fecal”, text = “fecal”, “metagenome”, “fecal animal”, “pigs”. A summary of the studies used is provided in Table 1. Detailed information about the metagenomes retrieved from the databases is included in Supplementary Table S1.

**Table 1:** Summary of included studies, sample number for each group of pigs (AB+/AB-), antibiotic usage (usage), treatment and antibiotic type (type) used in the study.

| Pigs | Study | Sample no. | Usage | Treatment | Type | Citation |
| --- | --- | --- | --- | --- | --- | --- |
| AB+ | 1 | 10 | Yes | Raised | Tetracycline | [14] |
| AB- | 2 | 27 | No | - |  | [15] |
| AB+ | 3 | 11 | Yes | Raised | Antifolates, tetracycline, beta-lactams | [16] |
| AB- |  | 9 | Yes | Illness | Antifolates, chloramphenicol, tetracycline, beta-lactams | [16] |
| AB- | 4 | 7 | No | - |  | [17] |
| AB+ | 5 | 26 | Yes | Raised | Oxytetracycline | [18] |
| AB- |  | 13 | No | - |  | [18] |

### 1.2 Bioinformatic analysis

To ensure that good quality reads were used throughout the study, quality control of reads was done using FastQC v.0.11.9 [19], and filtered using Trimmomatic v.0.39 [20]. Reads were trimmed for quality and adaptor contamination using Trimmomatic [20] while specifying ILLUMINACLIP: adapters.fa:2:30:10 LEADING:8 TRAILING:8 MINLEN:30 SLIDINGWINDOW: 4:20 parameters. The microbiome of pigs AB- and AB+ groups were profiled by Kraken2 v2.1.0 [21] using the minikraken2 microbial database (2019, 8GB). Thereafter, species confirmation and the estimation of abundance were done by Bracken v2.6.2 [22]. The relative abundances of bacterial genera and species were used to determine the composition of the microbial population. The average relative abundances of the bacterial species were used to calculate the alpha and beta-diversity between the groups of pigs raised with and without antibiotics.

Principal component analysis (PCA) was performed in R v.2.6-4 using the prcomp function to determine the differences between fecal samples obtained from pigs raised with and without antibiotics. Scores for each principal component (PC1 and PC2) of the microbiome were plotted on the x-axis and y-axis. The level of correlation was observed at the species level.

For the alpha-diversity, the Shannon diversity index was used to determine the diversity of the microbiome and resistome using the PAST v.4.03 software [23]. The Kolmogorov-Smirnov test was used to assess whether there were significant differences in the Shannon diversity indices between the AB+ and AB- groups using the scipy.stats package in Python. A p-value greater than 0.05 (p *≥* 0.05) indicated significant differences in the microbiome and resistome.

The beta-diversity with the Bray-Curtis dissimilarity was used to assess the difference between the fecal microbiome and resistome of pigs raised with and without antibiotics. The beta-diversity was visualised using Non-Metric Multidimensional Scaling (NMDS) and was conducted using the vegan package in R v.2.6-4. The goodness of fit for the NMDS ordination was assessed with 999 permutations to determine the significance of the clustering patterns using the envfit function in the vegan package in R. Differences in the beta-diversity between pigs AB+ and AB- were considered significant at p *≤* 0.05.

The filtered reads were *de novo* assembled using MegaHit v.1.2.9 [24] with default parameters. Assembled metagenomes were annotated to determine the antibiotic resistance gene families and their corresponding drug classes using the Comprehensive Antimicrobial Resistance Database Resistance Gene Identifier (CARD-RGI) v.3.1.2 with default settings including loose matches and the –low quality –clean options [25]. The version of the database was kept consistent throughout the analysis. The relative abundances of the annotated antibiotic resistance genes in each group (AB+ and AB-) were assessed. This data was normalised based on the total number of antibiotic resistance genes detected in each group, indicating their respective proportions. The covariance between the bacterial species and antibiotic resistance genes was calculated in Python v.3.7 using numpy v.1.23.4.

*De novo* metagenomic contigs were aligned for the detection of integrons and integrative elements by alignment to the INTEGRALL database [26] and the ICEberg database [27]. BWA-mem v.0.7.10 with default parameters was used to perform this alignment [28], generating a SAM file. To filter soft and hard clipped reads and undesirable alignments, the SAM files were filtered by the Samclip tool with default parameters. The output SAM file was converted into a BAM file using SAMtools v.1.9 [29]. Contigs that aligned to the databases were considered integrons and integrative conjugative elements.

PlasClass v.0.1.1 [30] was used to classify if plasmid-originating contigs. To ensure that contigs were truly plasmid-originating, a threshold of length *≥* 1000 and probability *≥* 0.75 was used. To determine if the detected plasmids were associated with the found antibiotic resistance genes, the plasmid-originating contigs were matched to the previously determined contigs carrying antibiotic resistance genes.

## 2 Results and Discussion

In this study, publicly available metagenomic data representing the fecal microbiome of pigs raised with (AB+) and without (AB-) antibiotics were analysed to identify and characterise the microbiome, resistome and mobilome.

### 2.1 The fecal microbiome is influenced by study-specific conditions and antibiotic usage

To investigate the effects of antibiotic usage on the taxonomic composition of the fecal microbiome between pigs raised with (AB+) and without (AB-) antibiotics, we conducted a PCA analysis using the average relative species abundances. Previous studies have shown that antibiotic administration in pigs alters the gut microbiome, consequently affecting the fecal microbiome of these animals [31].

In our study, five metagenomic datasets were used and categorised based on antibiotic usage (Table 1). In Figure 1 samples were colour-coded according to their studies with shapes representing antibiotic usage (Table 1). A smaller distance between samples on the PCA plot indicates a greater similarity between samples.

**Fig. 1:**
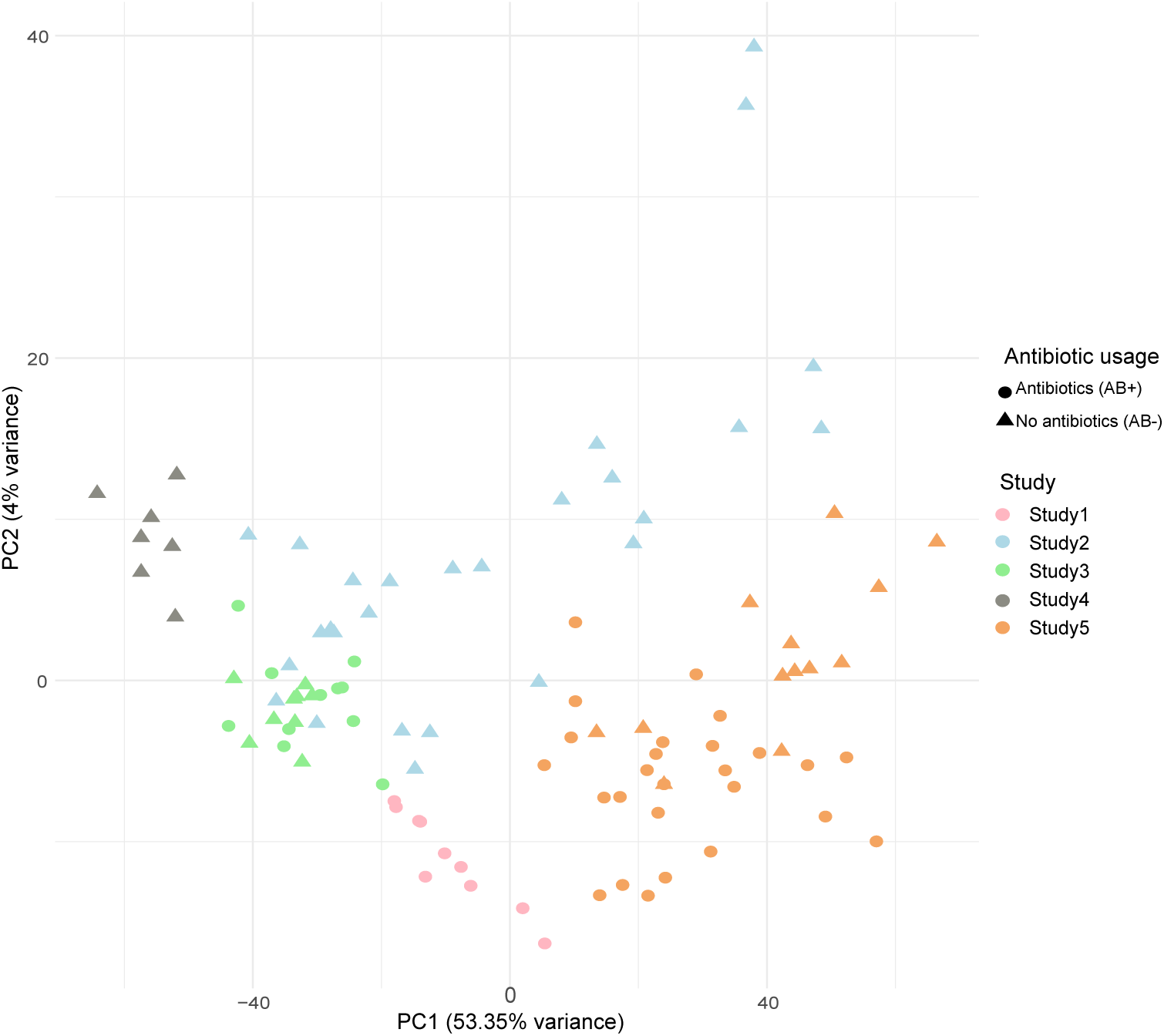
Visualisation of the PCA analysis using the average relative abundance of the bacterial species detected in fecal microbiome of pigs raised with (AB+) and without antibiotics (AB-) with the explained variance in brackets. Each colour represents a sample in a study (study 1= pink, study 2 = blue, study 3 = green, study 4 = grey, study 5 = orange) and shapes represent antibiotic usage (antibiotics or AB+ = circle and no antibiotics or AB- = triangle).

Antibiotic usage affects the fecal microbiome in combination with the study’s origin. Figure 1 shows the grouping of samples according to their respective studies. This grouping indicates that the microbial populations in each study are influenced by study-specific conditions. Specifically, samples grouped as AB+ and AB- obtained from study 3 and study 5 overlap within their respective studies. Study 3 sampled from Canada while Study 5 sampled from Austria. Unfortunately, due to the lack of metadata we cannot make inferences based on the study-specific conditions; however, these can include geographical location, pig diet and management practices. These factors can affect the composition of the fecal microbiome. When samples are labelled by antibiotic usage, as shown in Figure 1, samples group accordingly, indicating an impact on the composition of antibiotics on the fecal microbiome combined with study-specific conditions. Overall, pigs raised with and without antibiotic usage can have zoonotic pathogens such as *Clostridioides difficile* in the feces, as detected in our study (Figure 2) [32]. This can potentially spread to other environments such as soil, water, and other animals [33, 34].

**Fig. 2:**
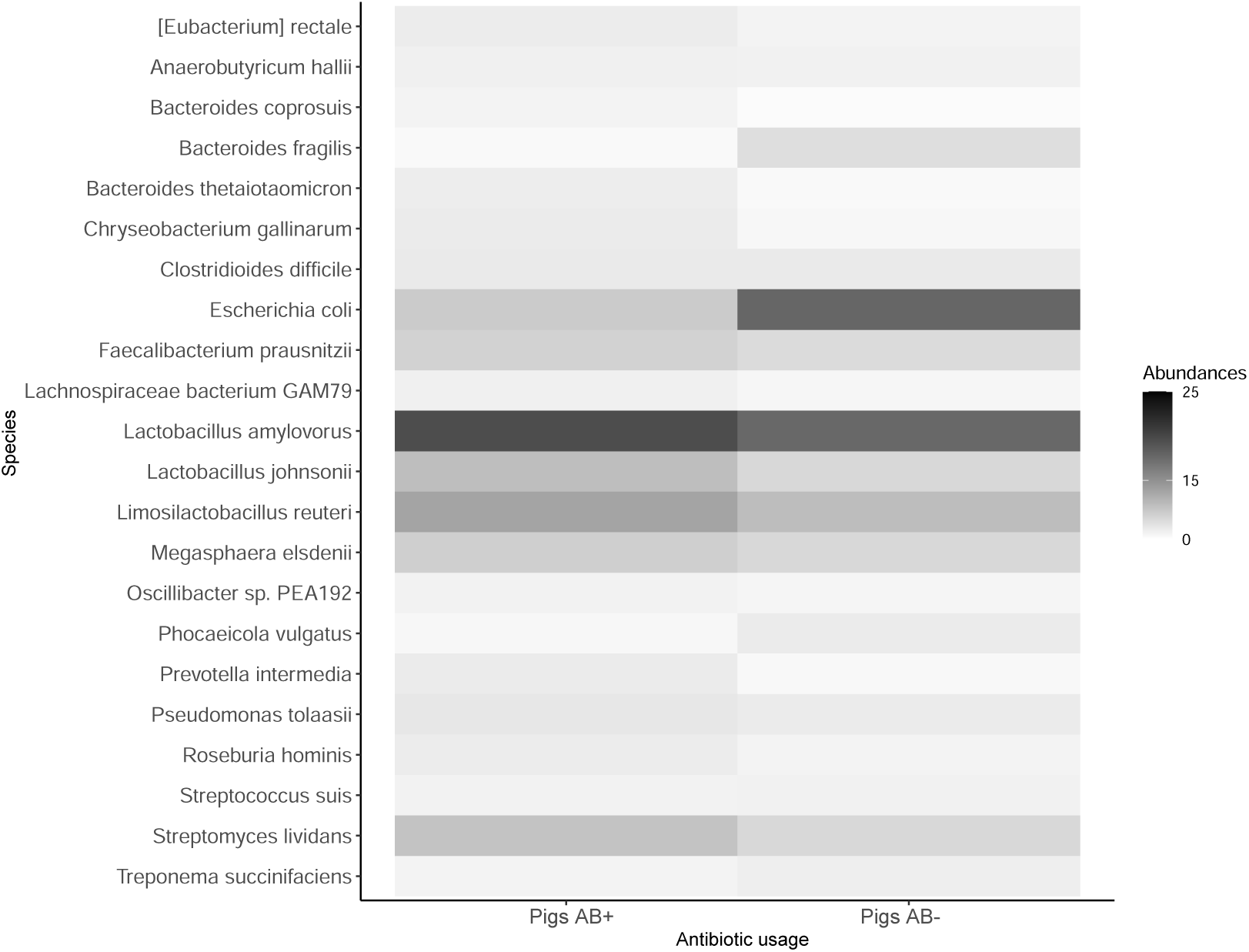
Heatmap showing the average relative abundance of bacterial species over 1% in the fecal microbiome, comparing pigs raised with (AB+) and without antibiotics (AB-). Higher abundances are indicated by the intensity of grey in the heatmap.

### 2.2 Probiotic bacteria are dominant in the fecal microbiome of pigs raised with and without antibiotics

To assess the differences in the microbial composition in the fecal microbiome of pigs AB+ and AB-, we performed taxonomic classification. Additionally, we investigated the similarities between the microbial communities to assess if antibiotic usage influences the bacterial species found and their abundances. Antibiotic administration has previously been shown to affect the indigenous microbes in animal feces leading to changes in the microbial community structure and resistance [35].

Based on the normalised data, a higher number of genera and species were present in the fecal microbiome of pigs raised without antibiotics (AB-). Specifically, we identified an total of 998 genera and 2580 species across the AB+ pigs studies compared to a total of 1024 genera and 2845 species across the AB- pigs studies. This increased number of species in the AB- group suggests that the absence of antibiotics allows for a more complex and varied microbial community (Supplementary Table S2-S3). It should be noted that the samples included were obtained from different studies therefore sample processing and sequencing depth can influence these results.

The higher number of bacterial species in pigs AB- may contribute to a more resilient gut ecosystem, potentially enhancing overall health and disease resistance [36]. 14 species that had an abundance of over 1% were in common between the two groups (Supplementary Table S5). This is supported by the Jaccard similarity of 0.685 indicating that 68% of the total bacteria present were shared between pigs AB+ and AB-. Additionally, the Shannon H diversity index was 4.018 for pigs AB+ compared to 4.007 for pigs AB- suggesting that pigs raised with antibiotics (AB+) have a more diverse microbiome as compared to pigs AB- (p *≥* 0.05). Details on the abundances of the bacterial genera, species, unique species and commonly identified species can be found in Supplementary Tables S2 - S5.

Probiotic bacteria are dominant in the fecal microbiome of pigs AB+ and AB- (Figure 2). We detected a higher average relative abundance of Lactobacillus genera (Supplementary Table S2), specifically *Lactobacillus amylovorus* in the fecal microbiome of pigs AB+ and AB-. The *Lactobacilli* species is one of the dominant bacterial groups establishing a stable population in the intestinal tract of piglets [37]. *Lactobacillus* is also a probiotic candidate known to have antibacterial properties. In animal husbandry, *Lactobacillus amylovorus* is a feed additive which assists in the daily weight gain of animals and resistance against enteric pathogens such as *Salmonella enterica* and *Escherichia coli* [38–40].

Combining probiotic bacteria with antibiotics may suppress opportunistic pathogens (Figure 2). In our study, we observed the *Escherichia* genera are abundant in the fecal microbiome of pigs AB- with most of the species attributed to the enteric opportunistic pathogen, *Escherichia coli* (*E. coli*). Contrastingly, this abundance is low in pigs AB+ indicating that the use of antibiotics in the growth of pigs may suppress the proliferation of *E. coli*. Previous studies have shown that while *E. coli* isolates are found in the feces of pigs before antibiotic administration, the use of antibiotics can suppress these opportunistic pathogens leading to changes in the abundances [41, 42]. Furthermore, we identified the unique bacterial species present in pigs AB+ and AB- (Supplementary Table S4). While we see a mixture of pathogenic bacteria and those that support gut health, pathogenic bacteria such as *Legionella pneumophila*, *Yersinia enterocolitica*, and *Vibrio cholerae* were identified in the feces of pigs AB+ and not in AB-. On the other hand, beneficial gut-supporting bacteria such as *Bifidobacterium* were identified in pigs AB- indicating a healthier gut microbiome which can lead to decreased diarrhoea incidence and the inhibition of pathogens [43].

As stated by Rochegue et al. [44], antibiotics usage may reduce microbial diversity however, this was not evident in our study even though pigs AB+ and AB- exhibited differences in the microbial community. The use of antibiotics can influence the microbial community and potentially promote the selection of antibiotic resistant bacteria.

#### 2.2.1 Variation in the microbial community of AB- pigs

To determine the differences in microbial communities found between the fecal microbiome of pigs AB+ and AB-, we measured the beta-diversity using the Bray-Curtis dissimilarity index. This index quantifies the distance between microbial communities based on species abundances. Visualised with non-metric multidimensional scaling (NMDS), a smaller distance between samples within a group i.e., AB+ or AB-, indicates a greater similarity between these microbial communities [45].

Figure 3 illustrates the Non-Metric Multidimensional Scaling (NMDS) analysis of the fecal microbiome, highlighting differences in the microbial populations between pigs raised with antibiotics (AB+) and those without antibiotics (AB-). Figure 3 shows the AB+ samples are more tightly grouped with less distance between them, indicating a high similarity in their microbial communities.

**Fig. 3:**
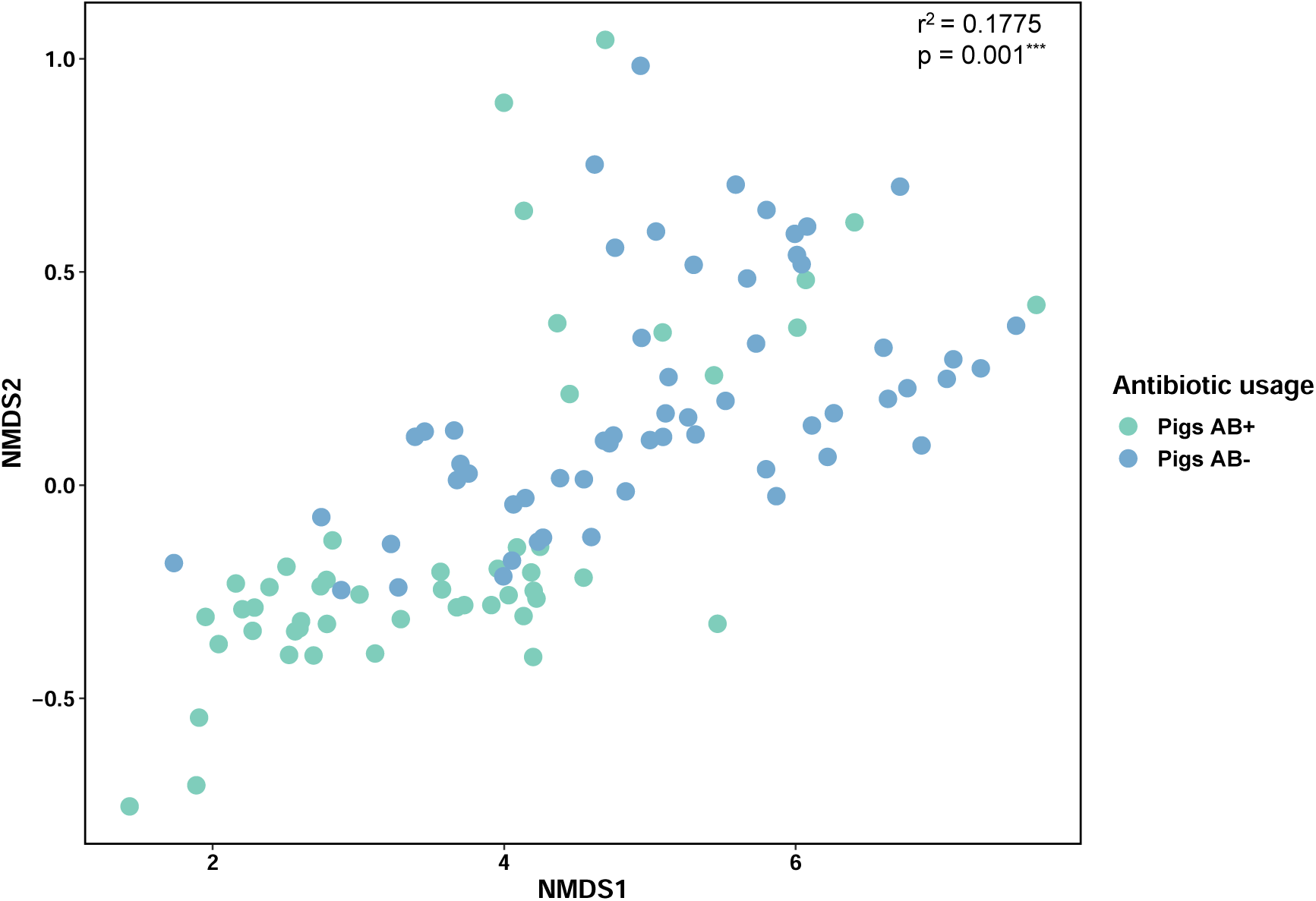
Visualisation of the beta-diversity analysis of the microbial community by Non-metric Multi-dimensional Scaling Ordination (NMDS) using Bray-Curtis dissimilarities between the fecal microbiome of pigs raised with and without antibiotics. Samples for each antibiotic usage group are indicated in different colours (Pigs AB+ = light blue, Pigs AB- = dark blue). Variation (*r*^2^) and p-value, calculated using the goodness of fit, is indicated in the top right corner with ^∗∗∗^ denoting the statistical significance of the difference between pigs AB+ and AB-.

In contrast, the AB- samples are more dispersed as shown by Figure 3, reflecting greater variability in their microbial communities. This implies that the microbiome of untreated pigs may be more influenced by external factors such as diet, environmental conditions, and other study-specific variables. The diversity in the AB- group indicates a more heterogeneous microbial population, potentially contributing to different health and functional outcomes such as a balanced immune system and better digestion [46]. Statistical analysis confirms the effect of antibiotics on microbial diversity, with antibiotics explaining 17.75% of the variation in microbial communities (r² = 0.1775, p = 0.001). Previous research has shown that less diversity in the gut microbiome of pigs is an indication of poor health. As stated previously, external factors can influence the gut microbiome however, microbial infections and the constant use of antibiotics lead to pigs being in an unhealthy state [47].

### 2.3 Similar AMR gene families are present in pigs raised with and without antibiotics

To investigate the effects of antibiotic usage on antibiotic resistance genes (ARGs) in the fecal microbiome of pigs AB+ and AB-, we identified the AMR gene families as bacteria that are constantly exposed to antibiotics may carry different ARGs [48]. Furthermore, we determined the differences in the AMR gene family composition by conducting a beta-diversity analysis visualised by NMDS. Similar to our analysis of the microbial composition, a smaller distance between samples of the AB+ or AB- groups indicates a greater similarity in AMR gene family composition. Additionally, we calculated the Shannon H diversity and Jaccard similarity index to assess the diversity within a group and if there are similar AMR gene families found between the pigs AB+ and AB- groups. Details can be found in Supplementary Tables S6 - S8.

Figure 4 shows the AMR gene families identified with an average relative abundance of over 1% in the fecal microbiome of pigs AB+ and AB-. The resistance nodulation cell division superfamily (RND) efflux pump, contributing to multi-drug resistance by expelling antibiotics from the bacterial cell wall, comprised 11% abundance of the AMR gene families detected in pigs AB- and 15% in pigs AB+ (Supplementary Table S6) [49]. Previous studies have shown the presence of RND efflux pump gene families is common in pig raised with and without antibiotics. Additionally, RND efflux pumps are commonly found in gram-negative bacteria e.g. *E. coli* and can export *β*-lactams out of the cell membrane [16, 50–53]. Interestingly, in our study, the presence of *E. coli* combined with the use of *β*–lactam antibiotics in feed (AB+) and to treat illness (AB-) may have contributed to the high abundance of the RND efflux pump gene family observed.

**Fig. 4:**
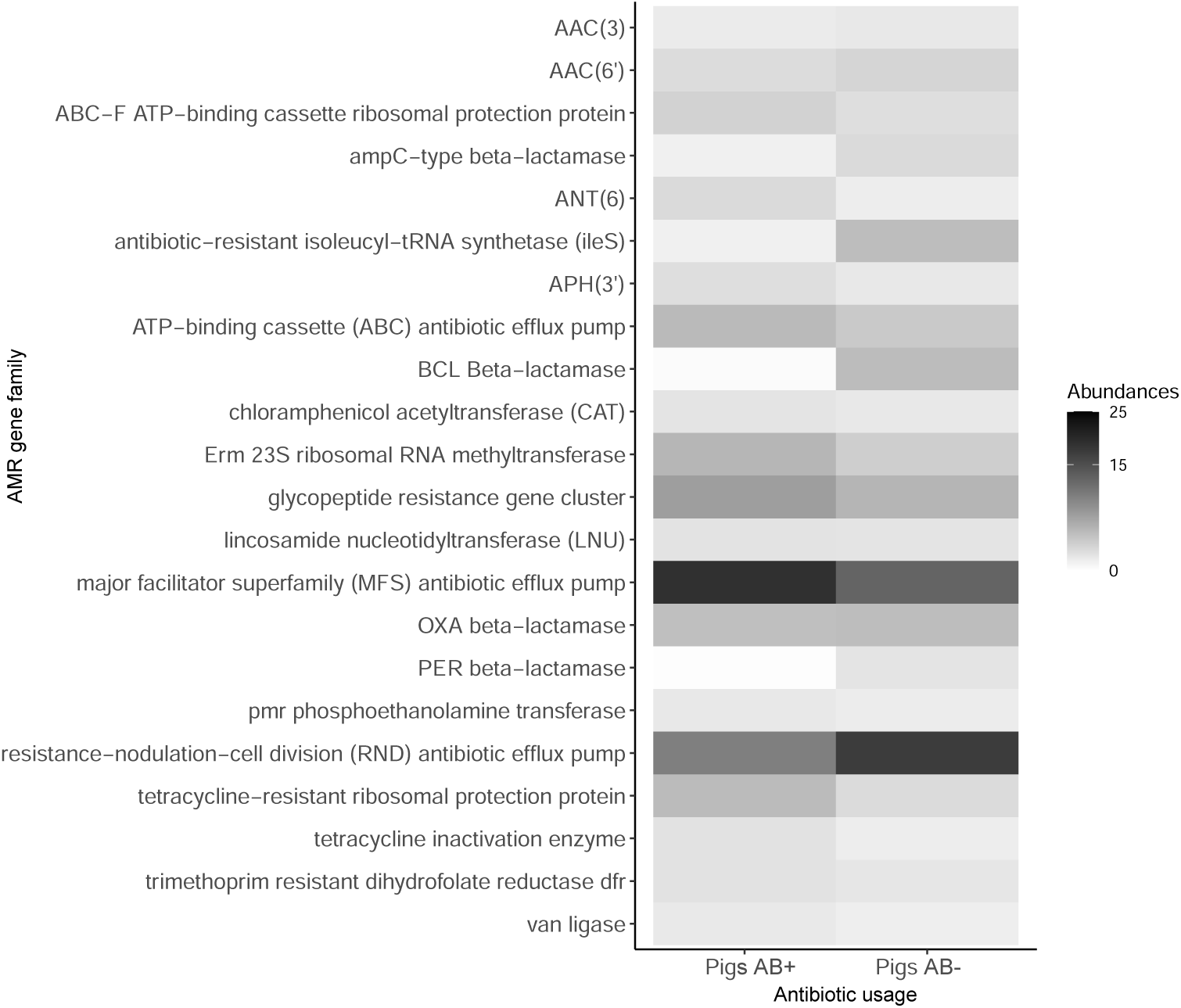
Heatmap showing the average relative abundance of ARGs over 1% in the fecal microbiome, comparing pigs raised with (AB+) and without antibiotics (AB-). Higher abundances are indicated by the intensity of grey in the heatmap.

Additionally, the OXA *β*-lactamase gene family had similar average relative abundance among the AMR gene families detected in the fecal microbiome of AB- pigs (3%) as compared to AB+ (3.5%) (Figure 4 and Supplementary Table S6). Since *β*-lactams are the most sold antibiotics used for food-producing animals i.e., pigs, it was expected that there would be a higher abundance of the OXA *β*-lactamase gene family present in AB+ samples. However, pigs raised without antibiotics or organically can be more exposed to environmental sources, pathogens, and ARGs [54]. The abundance of the OXA *β*-lactamase gene family in pigs AB- could also be attributed to either the administration of *β*-lactams to treat illnesses or injuries, horizontal gene transfer, management practices in farms or environmental differences [55, 56].

Antibiotic usage causes variation in AMR gene families detected. Figure 5 illustrates the beta-diversity of AMR gene families present in pigs treated with antibiotics (AB+) and those not treated with antibiotics (AB-). Samples within the AB+ group are tightly clustered together whereas samples from the AB- group have larger distances between them indicating variation in the resistome between both groups. This is further supported by the Shannon H diversity indices of 3.968 for AB+ and 4.052 for AB- (p*≥*0.05). This variation in AMR gene families can be caused by antibiotic usage (r² = 0.1768, p = 0.001). Interestingly, variations in AMR gene families are similar to the variation reported previously with the microbial communities (r² = 0.1775, p = 0.001) however, covariation analysis has shown that minimal co-variation occurs between the bacteria and ARGs identified suggesting that antibiotic usage is responsible for differences in AMR gene families and microbiome composition (Supplementary Table S7).

**Fig. 5:**
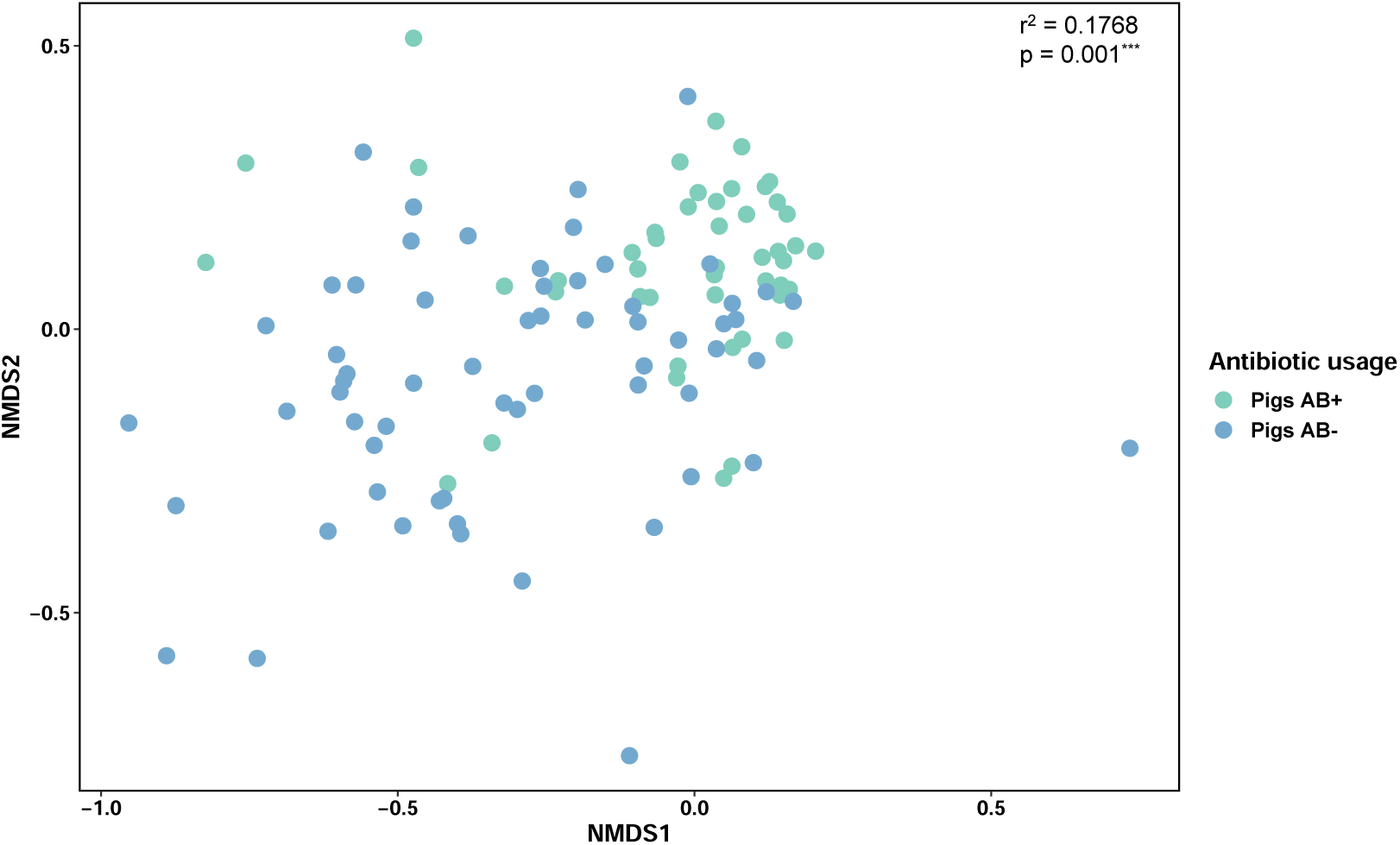
Visualisation of the beta-diversity of ARGs in the fecal microbiome of pigs raised with and without antibiotics by Non-Metric Multidimensional Scaling (NMDS). Samples are colour-coded based on antibiotic usage (Pigs AB+ = light blue, Pigs AB- = dark blue). Variation (*r*^2^) and p-value is indicated in the top right corner with ^∗∗∗^ denoting the statistical significance of the difference between pigs AB+ and AB-

Additionally, the Jaccard similarity index of 0.744 suggests that only 74% of the AMR gene families are shared between the AB+ and AB- groups, indicating an overlap in the AMR gene family composition between both groups. This can be seen in Figure 5, as some samples from the AB+ and AB- are close together while others are dispersed indicative of variation of AMR gene family composition. 11 AMR gene families present at over 1% were common between both groups (Supplementary Table S8). Differences in AMR gene families may be influenced by environmental factors, housing, or farm management practices, though more research is needed to fully understand the role of animal husbandry in shaping ARG distributions [56–58].

### 2.4 Plasmids linked to commonly used antibiotics

To investigate the presence of plasmids and their connection to antibiotic resistance in the fecal microbiome of pigs AB+ and AB-, we determined the percentage of contigs classified as plasmids. Furthermore, we examined their association with AMR gene families. Plasmids play a significant role in the spread of ARGs and AMR as bacteria and antibiotic residues present in feces can enhance horizontal gene transfer [9, 59, 60].

Over 1% of contigs were classified as plasmids in both groups. Further analysis showed that the AMR gene families were associated with 0.1% of the classified plasmids. Plasmids associated with AMR gene families were detected in both pigs AB+ and AB- (Figure 6 and Supplementary Table S9). In Figure 6 we observed a high average relative abundance of plasmids associated with the tetracycline-resistant ribosomal protection protein conferring resistance to the tetracycline antibiotic and the OXA *β*-lactamase gene family conferring multi-drug resistance to carbapenem; cephalosporin; penam.

**Fig. 6:**
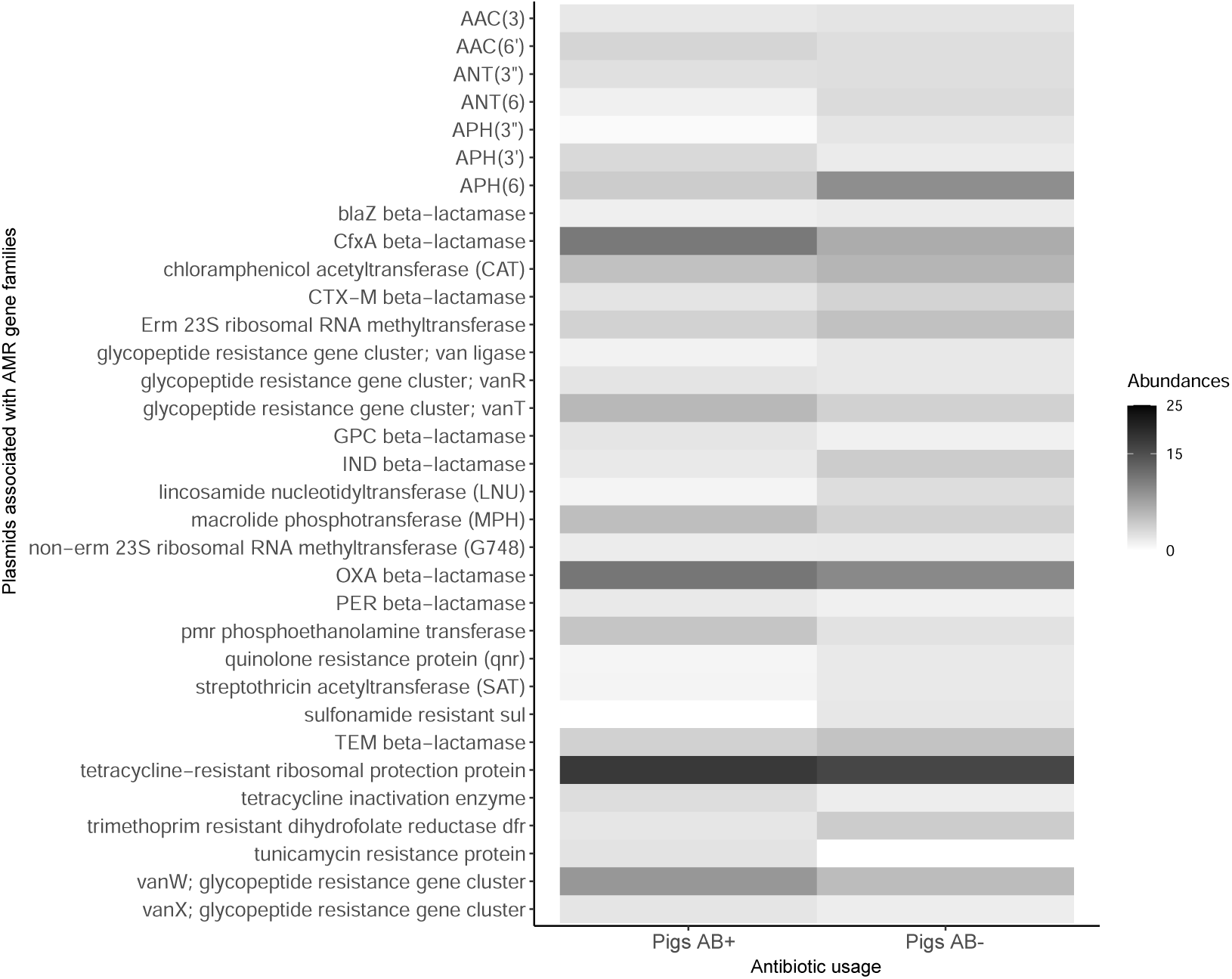
Heatmap showing the average relative abundances of plasmids associated with AMR gene families over 1% in the fecal microbiome, comparing pigs raised with (AB+) and without antibiotics (AB-). Higher abundances are indicated by the intensity of grey in the heatmap.

Plasmids linked to tetracycline resistance were detected in the fecal microbiome of pigs AB+ and AB- (11%). Tetracycline has been commonly used in animal feed and for the treatment of illnesses, including this study (Table 1 and Figure 6) [61]. Previous studies have discussed the continuous usage of tetracycline antibiotics selecting for plasmids associated with tetracycline resistance [8]. Furthermore, tetracycline resistance genes have been detected in soil that has been amended with animal manure indicating the transfer of genes via plasmids from one environment to another [62–65].

Secondly, plasmids associated with the OXA *β*-lactamase gene family were present at an abundance of 6-7% of the AMR-associated plasmids detected in pigs AB+ and AB- (Figure 6 and Supplementary Table S9). The OXA *β*-lactamase gene family confers multi-drug resistance to carbapenem; cephalosporin and penams. Similar to the use of tetracycline, *β*-lactams were used in both groups as feed additives or for the treatment of illnesses [18]. Plasmids associated with *β*-lactams are a public health concern as these can be transmitted through environmental sources such as manure, soil, air, wildlife and humans, especially farm workers [66, 67].

The detection of plasmids associated with tetracycline resistance and OXA *β*-lactams is not surprising as previous research has indicated that piglets can exhibit a pre-existing resistance pattern from birth that reflects the environment [68]. This aligns with the data used in our study, as antibiotics such as tetracycline, oxytetracycline, and beta-lactams were used in pigs raised with antibiotics. The use of antibiotics even for treatment can enhance the selection for ARGs and the horizontal transfer of these genes by plasmids. While the use of antibiotics is a common practice for therapeutic purposes, the low dosages can contribute to the emergence of antibiotic resistant bacteria thereby spreading to animals and humans [18, 31].

### 2.5 Class 1 integrons linked to chloramphenicol resistance

To investigate the presence of integrons in the fecal microbiome of pigs AB+ and AB-, we classified contigs as integrons. Integrons are responsible for the spread of AMR via horizontal gene transfer between pathogenic and non-pathogenic bacteria found in all environments [69]. In our study, we determined the number of integrons present in the contigs from pigs AB+ and AB-. We then classified them into classes 1, 2, and 3 to determine their abundances. Furthermore, we linked the detected integrons to ARGs to assess their role in AMR dissemination (Supplementary Table S10).

Firstly, less than 0.01% of contigs were classified as integrons. From the detected integrons we classified them into their respective groups of classes 1, 2 and 3. Table 2 shows that class 1 integrons were more abundant as compared to classes 2 and 3. Class 1 and 3 integrons are commonly found in freshwater and soil environments whereas class 2 integrons are found in marine environments [69]. Additionally, bacteria bearing class 1 integrons have been reported in environmental reservoirs such as farms, agricultural land and wastewater systems [70, 71]. It has been stated that the abundance of class 1 integrons can be identified in millions to billions of copies in a single gram of feces from agricultural animals [72].

**Table 2:** The average relative abundances of integrons in their respective classes detected in the fecal microbiome of pigs raised with (AB+) and without antibiotics (AB-).

| Classes of Integrons | Pigs AB+ | Pigs AB- |
| --- | --- | --- |
| Class 1 | 18% | 14% |
| Class 2 | 5% | 3% |
| Class 3 | 1% | 1% |

Class 1 integrons associated with the *cmxA* antibiotic resistance gene was detected in a higher abundance in pigs AB+ (30%) (Supplementary Table S10). This abundance was lower in AB- (15%). Chloramphenicol has been used to treat illness in our study, however, we have no information on the use of chloramphenicol in pigs raised with antibiotics. Previous research has shown that *cmxA* gene is linked to chloramphenicol resistance and has been detected in pig feces and in amended agricultural soils [70, 71, 73, 74]. This indicates that these ARGs can spread from livestock farms and animal feces to pathogens in soils via horizontal gene transfer i.e. integrons [73, 74].

A similar trend was observed with class 3 integrons as 71% of the class 3 integrons detected in pigs AB+ were associated with *blaOXA-2* gene. This was lower in the pigs AB- (39%) (Supplementary Table S9). This trend could be indicative of the selective pressure exerted by antibiotic use, where the presence of antibiotics may encourage the proliferation of resistant genes like *blaOXA-2* within integrons. Class 3 integrons are known for their role in antibiotic resistance and can capture and express genes conferring resistance to various antibiotics such as *β*-lactams [75, 76].

The prevalence of such genes associated with integrons in livestock is a concern for public health, as it may contribute to the spread of antimicrobial resistance from animals to humans through the food chain or other means of zoonotic transfer. It’s important to monitor these trends and implement strategies to mitigate the spread of resistance genes, ensuring both animal health and public safety [77].

### 2.6 Integrative elements detected in similar abundances

To identify the presence of integrative elements in the fecal microbiome of pigs AB+ and AB-, we determined the proportion (%) of integrative elements present in the contigs. Integrative elements are comprised of integrative and conjugative elements (ICEs), integrative and mobilisable elements (IMEs) and *cis*-integrative and mobilisable elements (CIMEs).

Integrative elements were detected in less than 0.0001% of contigs in both pigs AB+ and AB-. From those detected, approximately 50% of the integrative elements were ICEs, similar to a previous study [78]. ICE families were identified in both antibiotic groups, ie, AB + and AB- (Supplementary Table S11). Tn916 made up 2-3% of the detected ICE families and has been classified as a conjugative transposon which can carry ARGs. Tn916 has also been linked to pathogenic bacteria such as *Enterococcus fecalis* and *Staphylococcus aureus* [79]. This highlights the potential risk of AMR spreading within and between bacterial populations in livestock which can have implications for animal health and food safety.

The high abundance of various transposons among the identified IME families further emphasises the dynamic nature of genetic elements within these bacterial communities (Supplementary Table S12). Transposons are mobile genetic elements that can move within and between genomes, contributing to genetic diversity and adaptability [80]. Their prevalence in both AB+ and AB- groups suggests a widespread potential for genetic exchange and the evolution of bacterial populations in response to environmental pressures, such as the use of antibiotics.

Overall, these findings underscore the importance of monitoring integrative elements and their associated families, such as ICEs and transposons, to better understand their impact on bacterial communities within livestock and the broader implications for antibiotic resistance management.

## 3 Conclusion

We conducted a comprehensive analysis of the fecal microbiome of pigs raised with (AB+) and without (AB-) antibiotics to assess how antibiotic exposure influences the microbial community, antibiotic resistance genes (ARGs), and mobile genetic elements (MGEs).

Our study demonstrated that antibiotic usage influences the fecal microbiome of pigs together with the farm and environmental conditions specific to the studies used. Notably, while both pigs AB+ and AB- exhibited a high average relative abundance of the probiotic bacteria *Lactobacilli*, there was a difference in the diversity of bacterial species between the two groups. The use of antibiotics contributes to the variation in the fecal microbiome of pigs, however, it may not strongly influence the overall microbial community structure.

Similarly, a diverse set of antibiotic resistance genes was observed in the AB- group, underscoring the impact of antibiotic use on the gut microbiome. However, this impact does not create a huge variation between the resistome of AB+ and AB- as indicated by the Shannon diversity index. Unfortunately, this study is limited in sample size and metadata to confirm the influence of the antibiotics on the microbial community, resistome and mobilome.

We also identified the widespread presence of the RND efflux pump gene family in both groups, although the OXA *β*-lactamase gene was more prevalent in the AB- group. Additionally, plasmids associated with tetracycline and OXA *β*-lactamase resistance were detected. Integrons carrying chloramphenicol resistance genes and integrative elements linked to transposons were also identified, further illustrating the complexity of ARG dissemination in these environments.

Understanding the impact of antibiotic usage on the gut microbiome of pigs is critical for comprehending the spread of ARGs, especially since a significant portion of ARGs and antibiotic-resistant bacteria are excreted in feces. Although our study provides valuable insights into how antibiotic use affects the fecal microbiome, it is limited by the sample size, the absence of metadata and the fact that fecal samples may not fully represent the entire gut microbiome. Nevertheless, these findings highlight the potential for fecal matter to influence environmental microbial communities and contribute to the broader dissemination of antibiotic resistance. Future research should focus on the broader sampling of the gut microbiome to better understand the full impact of antibiotic use in agricultural settings.

## Declarations

### 3.0.1 Funding

This work is based on the research supported wholly/in part by the National Research Foundation of South Africa (Grant Numbers: 120192).

### 3.0.2 Conflict of interest/Competing interests

The authors declare that the research was conducted in the absence of any commercial or financial relationships that could be construed as a potential conflict of interest.

### 3.0.3 Ethics approval and consent to participate

Not applicable

### 3.0.4 Consent for publication

Not applicable

### 3.0.5 Data availability

These datasets generated and/or analysed during the current study are available in the NCBI SRA repository.

- Study 1: https://www.ncbi.nlm.nih.gov/sra/PRJEB12458
- Study 2: https://www.ncbi.nlm.nih.gov/sra/PRJNA629856
- Study 3: https://www.ncbi.nlm.nih.gov/sra/PRJNA737271
- Study 4: https://www.ncbi.nlm.nih.gov/sra/PRJNA844176
- Study 5: https://www.ncbi.nlm.nih.gov/sra/PRJNA471402

Details on the SRA run numbers are listed in Supplementary Table 1.

### 3.0.6 Materials availability

Not applicable

### 3.0.7 Code availability

Not applicable

### 3.0.8 Author contribution

SP and TA designed experiments. SP conducted all experiments, analysed data and wrote the manuscript. SP and TA edited and proofread the manuscript. All authors contributed to the article and approved the submitted version.

## Appendix A Supplementary material

Supplementary information can be found in Supplementary material.xlsx

## Notes

### Competing Interest Statement

The authors have declared no competing interest.

## References

[1] Sivagami, K., Vignesh, V.J., Srinivasan, R., Divyapriya, G., Nambi, I.M.: Antibiotic usage, residues and resistance genes from food animals to human and environment: An indian scenario. Journal of Environmental Chemical Engineering 8(1), 102221 (2020)

[2] Tiseo, K., Huber, L., Gilbert, M., Robinson, T.P., Van Boeckel, T.P.: Global trends in antimicrobial use in food animals from 2017 to 2030. Antibiotics 9(12), 918 (2020)

[3] Thacker, P.A.: Alternatives to antibiotics as growth promoters for use in swine production: a review. Journal of animal science and biotechnology 4, 1–12 (2013)

[4] Lekagul, A., Tangcharoensathien, V., Mills, A., Rushton, J., Yeung, S.: How antibiotics are used in pig farming: a mixed-methods study of pig farmers, feed mills and veterinarians in thailand. BMJ Global Health 5(2), 001918 (2020)

[5] Mencía-Ares, O., Cabrera-Rubio, R., Cobo-Díaz, J.F., Álvarez-Ordóñez, A., Gómez-García, M., Puente, H., Cotter, P.D., Crispie, F., Carvajal, A., Rubio, P., et al.: Antimicrobial use and production system shape the fecal, environmental, and slurry resistomes of pig farms. Microbiome 8(1), 1–17 (2020)

[6] Chan, R., Chiemchaisri, C., Chiemchaisri, W., Boonsoongnern, A., Tulayakul, P.: Occurrence of antibiotics in typical pig farming and its wastewater treatment in thailand. Emerging Contaminants 8, 21–29 (2022)

[7] Lekagul, A., Tangcharoensathien, V., Yeung, S.: Patterns of antibiotic use in global pig production: a systematic review. Veterinary and animal science 7, 100058 (2019)

[8] Leclercq, S.O., Wang, C., Zhu, Y., Wu, H., Du, X., Liu, Z., Feng, J.: Diversity of the tetracycline mobilome within a chinese pig manure sample. Applied and Environmental Microbiology 82(21), 6454–6462 (2016)

[9] Zalewska, M., Bl-ażejewska, A., Czapko, A., Popowska, M.: Pig manure treatment strategies for mitigating the spread of antibiotic resistance. Scientific Reports 13(1), 11999 (2023)

[10] Matheson, S., Edwards, S., Kyriazakis, I.: Farm characteristics affecting antibiotic consumption in pig farms in england. Porcine Health Management 8(1), 7 (2022)

[11] Hu, J., Chen, J., Ma, L., Hou, Q., Zhang, Y., Kong, X., Huang, X., Tang, Z., Wei, H., Wang, X., et al.: Characterizing core microbiota and regulatory functions of the pig gut microbiome. The ISME Journal 18(1), 037 (2024)

[12] Lemon, K.P., Armitage, G.C., Relman, D.A., Fischbach, M.A.: Microbiota-targeted therapies: an ecological perspective. Science translational medicine 4(137), 137–51375 (2012)

[13] Ahn, J.-S., Lkhagva, E., Jung, S., Kim, H.-J., Chung, H.-J., Hong, S.-T.: Fecal microbiome does not represent whole gut microbiome. Cellular Microbiology 2023(1), 6868417 (2023)

[14] Munk, P., Andersen, V.D., Knegt, L., Jensen, M.S., Knudsen, B.E., Lukjancenko, O., Mordhorst, H., Clasen, J., Agersø, Y., Folkesson, A., et al.: A sampling and metagenomic sequencing-based methodology for monitoring antimicrobial resistance in swine herds. Journal of Antimicrobial Chemotherapy 72(2), 385–392 (2017)

[15] Holman, D.B., Kommadath, A., Tingley, J.P., Abbott, D.W.: Novel insights into the pig gut microbiome using metagenome-assembled genomes. Microbiology Spectrum 10(4), 02380–22 (2022)

[16] Chekabab, S.M., Lawrence, J.R., Alvarado, A.C., Predicala, B.Z., Korber, D.R.: Piglet gut and in-barn manure from farms on a raised without antibiotics program display reduced antimicrobial resistance but an increased prevalence of pathogens. Antibiotics 10(10), 1152 (2021)

[17] Wei, L., Zhou, W., Zhu, Z.: Comparison of changes in gut microbiota in wild boars and domestic pigs using 16s rrna gene and metagenomics sequencing technologies. Animals 12(17), 2270 (2022)

[18] Ghanbari, M., Klose, V., Crispie, F., Cotter, P.D.: The dynamics of the antibiotic resistome in the feces of freshly weaned pigs following therapeutic administration of oxytetracycline. Scientific reports 9(1), 4062 (2019)

[19] Andrews, S., et al.: FastQC: a quality control tool for high throughput sequence data. Babraham Bioinformatics, Babraham Institute, Cambridge, United Kingdom (2010)

[20] Bolger, A.M., Lohse, M., Usadel, B.: Trimmomatic: a flexible trimmer for illumina sequence data. Bioinformatics 30(15), 2114–2120 (2014)

[21] Wood, D.E., Lu, J., Langmead, B.: Improved metagenomic analysis with kraken 2. Genome biology 20(1), 1–13 (2019)

[22] Lu, J., Breitwieser, F.P., Thielen, P., Salzberg, S.L.: Bracken: estimating species abundance in metagenomics data. PeerJ Computer Science 3, 104 (2017)

[23] Hammer, Ø., Harper, D.A., Ryan, P.D., et al.: Past: Paleontological statistics software package for education and data analysis. Palaeontologia electronica 4(1), 9 (2001)

[24] Li, D., Luo, R., Liu, C.-M., Leung, C.-M., Ting, H.-F., Sadakane, K., Yamashita, H., Lam, T.-W.: Megahit v1. 0: a fast and scalable metagenome assembler driven by advanced methodologies and community practices. Methods 102, 3–11 (2016)

[25] McArthur, A.G., Waglechner, N., Nizam, F., Yan, A., Azad, M.A., Baylay, A.J., Bhullar, K., Canova, M.J., De Pascale, G., Ejim, L., et al.: The comprehensive antibiotic resistance database. Antimicrobial agents and chemotherapy 57(7), 3348–3357 (2013)

[26] Moura, A., Soares, M., Pereira, C., Leitão, N., Henriques, I., Correia, A.: Integrall: a database and search engine for integrons, integrases and gene cassettes. Bioinformatics 25(8), 1096–1098 (2009)

[27] Bi, D., Xu, Z., Harrison, E.M., Tai, C., Wei, Y., He, X., Jia, S., Deng, Z., Rajakumar, K., Ou, H.-Y.: Iceberg: a web-based resource for integrative and conjugative elements found in bacteria. Nucleic acids research 40(D1), 621–626 (2012)

[28] Li, H.: Aligning sequence reads, clone sequences and assembly contigs with bwamem. arXiv preprint arXiv:1303.3997 (2013)

[29] Li, H., Handsaker, B., Wysoker, A., Fennell, T., Ruan, J., Homer, N., Marth, G., Abecasis, G., Durbin, R.: The sequence alignment/map format and samtools. Bioinformatics 25(16), 2078–2079 (2009)

[30] Pellow, D., Mizrahi, I., Shamir, R.: Plasclass improves plasmid sequence classification. PLoS computational biology 16(4), 1007781 (2020)

[31] Ricker, N., Trachsel, J., Colgan, P., Jones, J., Choi, J., Lee, J., Coetzee, J.F., Howe, A., Brockmeier, S.L., Loving, C.L., et al.: Toward antibiotic stewardship: route of antibiotic administration impacts the microbiota and resistance gene diversity in swine feces. Frontiers in veterinary science 7, 255 (2020)

[32] Rahman, M.T., Sobur, M.A., Islam, M.S., Ievy, S., Hossain, M.J., El Zowalaty, M.E., Rahman, A.T., Ashour, H.M.: Zoonotic diseases: etiology, impact, and control. Microorganisms 8(9), 1405 (2020)

[33] Tang, X., Zhang, K., Xiong, K.: Fecal microbial changes in response to finishing pigs directly fed with fermented feed. Frontiers in Veterinary Science 9, 894909 (2022)

[34] Alvarado, A.C., Chekabab, S.M., Predicala, B.Z., Korber, D.R.: Impact of raised without antibiotics measures on antimicrobial resistance and prevalence of pathogens in sow barns. Antibiotics 11(9), 1221 (2022)

[35] Xiong, W., Wang, Y., Sun, Y., Ma, L., Zeng, Q., Jiang, X., Li, A., Zeng, Z., Zhang, T.: Antibiotic-mediated changes in the fecal microbiome of broiler chickens define the incidence of antibiotic resistance genes. Microbiome 6, 1–11 (2018)

[36] Guitart-Matas, J., Ballester, M., Fraile, L., Darwich, L., Giler-Baquerizo, N., Tarres, J., López-Soria, S., Ramayo-Caldas, Y., Migura-Garcia, L.: Gut microbiome and resistome characterization of pigs treated with commonly used post-weaning diarrhea treatments. Animal Microbiome 6(1), 24 (2024)

[37] Marti, R., Dabert, P., Ziebal, C., Pourcher, A.-M.: Evaluation of lactobacillus sobrius/l. amylovorus as a new microbial marker of pig manure. Applied and Environmental Microbiology 76(5), 1456–1461 (2010)

[38] Shen, J., Zhang, J., Zhao, Y., Lin, Z., Ji, L., Ma, X.: Tibetan pig-derived probiotic lactobacillus amylovorus slzx20-1 improved intestinal function via producing enzymes and regulating intestinal microflora. Frontiers in Nutrition 9, 846991 (2022)

[39] Petsuriyawong, B., Khunajakr, N.: Screening of probiotic lactic acid bacteria from piglet feces. Agriculture and Natural Resources 45(2), 245–253 (2011)

[40] Sirichokchatchawan, W., Pupa, P., Praechansri, P., Am-In, N., Tanasupawat, S., Sonthayanon, P., Prapasarakul, N.: Autochthonous lactic acid bacteria isolated from pig faeces in thailand show probiotic properties and antibacterial activity against enteric pathogenic bacteria. Microbial pathogenesis 119, 208–215 (2018)

[41] Cameron-Veas, K., Solá-Ginés, M., Moreno, M.A., Fraile, L., Migura-Garcia, L.: Impact of the use of *β*-lactam antimicrobials on the emergence of escherichia coli isolates resistant to cephalosporins under standard pig-rearing conditions. Applied and Environmental Microbiology 81(5), 1782–1787 (2015)

[42] Unno, T., Kim, J., Guevarra, R.B., Nguyen, S.G.: Effects of antibiotic growth promoter and characterization of ecological succession in swine gut microbiota. Journal of microbiology and biotechnology 25(4), 431–438 (2015)

[43] Pang, J., Liu, Y., Kang, L., Ye, H., Zang, J., Wang, J., Han, D.: Bifidobacterium animalis promotes the growth of weaning piglets by improving intestinal development, enhancing antioxidant capacity, and modulating gut microbiota. Applied and Environmental Microbiology 88(22), 01296–22 (2022)

[44] Rochegüe, T., Haenni, M., Mondot, S., Astruc, C., Cazeau, G., Ferry, T., Madec, J.-Y., Lupo, A.: Impact of antibiotic therapies on resistance genes dynamic and composition of the animal gut microbiota. Animals 11(11), 3280 (2021)

[45] Xia, Y., Sun, J., Chen, D.-G., Xia, Y., Sun, J., Chen, D.-G.: Community diversity measures and calculations. Statistical analysis of microbiome data with R, 167– 190 (2018)

[46] Jo, H.E., Kwon, M.-S., Whon, T.W., Kim, D.W., Yun, M., Lee, J., Shin, M.-Y., Kim, S.-H., Choi, H.-J.: Alteration of gut microbiota after antibiotic exposure in finishing swine. Frontiers in Microbiology 12, 596002 (2021)

[47] Upadhaya, S.D., Kim, I.H.: Maintenance of gut microbiome stability for optimum intestinal health in pigs–a review. Journal of Animal Science and Biotechnology 13(1), 140 (2022)

[48] Tao, S., Chen, H., Li, N., Wang, T., Liang, W.: The spread of antibiotic resistance genes in vivo model. Canadian Journal of Infectious Diseases and Medical Microbiology 2022(1), 3348695 (2022)

[49] Colclough, A.L., Alav, I., Whittle, E.E., Pugh, H.L., Darby, E.M., Legood, S.W., McNeil, H.E., Blair, J.M.: Rnd efflux pumps in gram-negative bacteria; regulation, structure and role in antibiotic resistance. Future microbiology 15(2), 143–157 (2020)

[50] Begmatov, S., Beletsky, A.V., Gruzdev, E.V., Mardanov, A.V., Glukhova, L.B., Karnachuk, O.V., Ravin, N.V.: Distribution patterns of antibiotic resistance genes and their bacterial hosts in a manure lagoon of a large-scale swine finishing facility. Microorganisms 10(11), 2301 (2022)

[51] Suriyaphol, P., Chiu, J.K.H., Yimpring, N., Tunsagool, P., Mhuantong, W., Chuanchuen, R., Bessarab, I., Williams, R.B., Ong, R.T.-H., Suriyaphol, G.: Dynamics of the fecal microbiome and antimicrobial resistome in commercial piglets during the weaning period. Scientific Reports 11(1), 18091 (2021)

[52] Keum, G.B., Kim, E.S., Cho, J., Song, M., Oh, K.K., Cho, J.H., Kim, S., Kim, H., Kwak, J., Doo, H., et al.: Analysis of antibiotic resistance genes in pig feces during the weaning transition using whole metagenome shotgun sequencing. Journal of Animal Science and Technology 65(1), 175 (2023)

[53] Routh, M.D., Zalucki, Y., Su, C.-C., Long, F., Zhang, Q., Shafer, W.M., Edward, W.Y.: Efflux pumps of the resistance-nodulation-division family: A perspective of their structure, function and regulation in gram-negative bacteria. Advances in enzymology and related areas of molecular biology 77, 109 (2011)

[54] Gerzova, L., Babak, V., Sedlar, K., Faldynova, M., Videnska, P., Cejkova, D., Jensen, A.N., Denis, M., Kerouanton, A., Ricci, A., et al.: Characterization of antibiotic resistance gene abundance and microbiota composition in feces of organic and conventional pigs from four eu countries. PLoS One 10(7), 0132892 (2015)

[55] Gāliņa, D., Balins, A., Valdovska, A.: The prevalence and characterization of fecal extended-spectrum-beta-lactamase-producing escherichia coli isolated from pigs on farms of different sizes in latvia. Antibiotics 10(9), 1099 (2021)

[56] Libisch, B., Abdulkadir, S., Keresztény, T., Papp, P.P., Olasz, F., Fébel, H., Sándor, Z.J., Rasschaert, G., Lambrecht, E., Heyndrickx, M., et al.: Detection of acquired antibiotic resistance genes in domestic pig (sus scrofa) and common carp (cyprinus carpio) intestinal samples by metagenomics analyses in hungary. Antibiotics 11(10), 1441 (2022)

[57] Soundararajan, M., Marincola, G., Liong, O., Marciniak, T., Wencker, F.D., Hofmann, F., Schollenbruch, H., Kobusch, I., Linnemann, S., Wolf, S.A., et al.: Farming practice influences antimicrobial resistance burden of non-aureus staphylococci in pig husbandries. Microorganisms 11(1), 31 (2022)

[58] Ager, E.O., Carvalho, T., Silva, E.M., Ricke, S.C., Hite, J.L.: Global trends in antimicrobial resistance on organic and conventional farms. Scientific Reports 13(1), 22608 (2023)

[59] Yang, F., Han, B., Gu, Y., Zhang, K.: Swine liquid manure: a hotspot of mobile genetic elements and antibiotic resistance genes. Scientific Reports 10(1), 15037 (2020)

[60] Wu, S., Dalsgaard, A., Hammerum, A.M., Porsbo, L.J., Jensen, L.B.: Prevalence and characterization of plasmids carrying sulfonamide resistance genes among escherichia coli from pigs, pig carcasses and human. Acta Veterinaria Scandinavica 52, 1–7 (2010)

[61] Granados-Chinchilla, F., Rodríguez, C.: Tetracyclines in food and feedingstuffs: from regulation to analytical methods, bacterial resistance, and environmental and health implications. Journal of analytical methods in chemistry 2017(1), 1315497 (2017)

[62] Schmitt, H., Stoob, K., Hamscher, G., Smit, E., Seinen, W.: Tetracyclines and tetracycline resistance in agricultural soils: microcosm and field studies. Microbial ecology 51, 267–276 (2006)

[63] Kazimierczak, K.A., Scott, K.P., Kelly, D., Aminov, R.I.: Tetracycline resistome of the organic pig gut. Applied and environmental microbiology 75(6), 1717–1722 (2009)

[64] Tams, K.W., Larsen, I., Hansen, J.E., Spiegelhauer, H., Strøm-Hansen, A.D., Rasmussen, S., Ingham, A.C., Kalmar, L., Kean, I.R.L., Angen, Ø., et al.: The effects of antibiotic use on the dynamics of the microbiome and resistome in pigs. Animal Microbiome 5(1), 39 (2023)

[65] Ma, X., Yang, Z., Xu, T., Qian, M., Jiang, X., Zhan, X., Han, X.: Chlortetracycline alters microbiota of gut or faeces in pigs and leads to accumulation and migration of antibiotic resistance genes. Science of the Total Environment 796, 148976 (2021)

[66] Cardoso, O., Osório, S., Ramos, F., Donato, M.M.: Plasmid-encoded ampc and extended-spectrum beta-lactamases in multidrug-resistant escherichia coli isolated from piglets in portugal. Microbial Drug Resistance 27(12), 1742–1749 (2021)

[67] Furlan, J.P.R., Stehling, E.G.: Detection of *β*-lactamase encoding genes in feces, soil and water from a brazilian pig farm. Environmental monitoring and assessment 190, 1–7 (2018)

[68] Scicchitano, D., Leuzzi, D., Babbi, G., Palladino, G., Turroni, S., Laczny, C.C., Wilmes, P., Correa, F., Leekitcharoenphon, P., Savojardo, C., et al.: Dispersion of antimicrobial resistant bacteria in pig farms and in the surrounding environment. Animal Microbiome 6(1), 17 (2024)

[69] Gillings, M.R.: Integrons: past, present, and future. Microbiology and molecular biology reviews 78(2), 257–277 (2014)

[70] Gaze, W.H., Zhang, L., Abdouslam, N.A., Hawkey, P.M., Calvo-Bado, L., Royle, J., Brown, H., Davis, S., Kay, P., Boxall, A.B., et al.: Impacts of anthropogenic activity on the ecology of class 1 integrons and integron-associated genes in the environment. The ISME journal 5(8), 1253–1261 (2011)

[71] Ye, C., Hou, F., Xu, D., Huang, Q., Chen, X., Zeng, Z., Peng, Y., Fang, R.: Prevalence and characterisation of class 1 and 2 integrons in multi-drug resistant isolates from pig farms in chongqing, china. Journal of Veterinary Research 64(3), 381–386 (2020)

[72] Ghaly, T.M., Gillings, M.R., Penesyan, A., Qi, Q., Rajabal, V., Tetu, S.G.: The natural history of integrons. Microorganisms 9(11), 2212 (2021)

[73] Zhou, B., Wang, C., Zhao, Q., Wang, Y., Huo, M., Wang, J., Wang, S.: Prevalence and dissemination of antibiotic resistance genes and coselection of heavy metals in chinese dairy farms. Journal of hazardous materials 320, 10–17 (2016)

[74] Xu, M., Wang, F., Sheng, H., Stedtfeld, R.D., Li, Z., Hashsham, S.A., Jiang, X., Tiedje, J.M.: Does anaerobic condition play a more positive role in dissipation of antibiotic resistance genes in soil? Science of The Total Environment 757, 143737 (2021)

[75] Deng, Y., Bao, X., Ji, L., Chen, L., Liu, J., Miao, J., Chen, D., Bian, H., Li, Y., Yu, G.: Resistance integrons: class 1, 2 and 3 integrons. Annals of clinical microbiology and antimicrobials 14, 1–11 (2015)

[76] Aruhomukama, D., Najjuka, C.F., Kajumbula, H., Okee, M., Mboowa, G., Sserwadda, I., Mayanja, R., Joloba, M.L., Kateete, D.P.: Bla vim-and bla oxa-mediated carbapenem resistance among acinetobacter baumannii and pseudomonas aeruginosa isolates from the mulago hospital intensive care unit in kampala, uganda. BMC infectious diseases 19, 1–8 (2019)

[77] Ejaz, H., Younas, S., Abosalif, K.O., Junaid, K., Alzahrani, B., Alsrhani, A., Abdalla, A.E., Ullah, M.I., Qamar, M.U., Hamam, S.S.: Molecular analysis of bla shv, bla tem, and bla ctx-m in extended-spectrum *β*-lactamase producing enterobacteriaceae recovered from fecal specimens of animals. PLoS One 16(1), 0245126 (2021)

[78] Wang, C., Song, Y., Tang, N., Zhang, G., Leclercq, S.O., Feng, J.: The shared resistome of human and pig microbiota is mobilized by distinct genetic elements. Applied and environmental microbiology 87(5), 01910–20 (2021)

[79] Roberts, A.P., Mullany, P.: Tn 916-like genetic elements: a diverse group of modular mobile elements conferring antibiotic resistance. FEMS microbiology reviews 35(5), 856–871 (2011)

[80] Chénais, B., Caruso, A., Hiard, S., Casse, N.: The impact of transposable elements on eukaryotic genomes: from genome size increase to genetic adaptation to stressful environments. Gene 509(1), 7–15 (2012)

